# *Gpr88* deletion impacts motivational control independently of striatal dopamine function

**DOI:** 10.1101/2022.05.19.492565

**Authors:** Daisy L. Spark, Michela H. Vermeulen, Patricia Rueda, Rocío de la Fuente Gonzalez, Tara Sepehrizadeh, Michael De Veer, Clotilde Mannoury la Cour, Alex Fornito, Monica Langiu, Gregory D. Stewart, Jess Nithianantharajah, Christopher J. Langmead

**Affiliations:** Drug Discovery Biology, Monash University, 381 Royal Parade, Parkville, VIC 3052, Australia; Neuroscience & Mental Health Therapeutic Program Area, Monash University, 381 Royal Parade, Parkville, VIC 3052, Australia; Neuromedicines Discovery Centre, Monash Institute of Pharmaceutical Sciences, Monash University, 381 Royal Parade, Parkville, VIC 3052, Australia; Monash Biomedical Imaging, Monash University, Clayton, VIC 3800, Australia; Institut de Recherches Servier, 50 Rue Carnot, 92150 Suresnes, France; Turner Institute for Brain and Mental Health, Monash Biomedical Imaging, and School of Psychological Sciences, Monash University Clayton, VIC, 3800, Australia; Florey Institute of Neuroscience and Mental Health, University of Melbourne, Parkville, VIC 3052, Australia

**Keywords:** GPR88, orphan GPCR, motivation, striatum, dopamine, touchscreen

## Abstract

**Background:** Disrupted motivational control is a common—but poorly treated—feature of psychiatric disorders. Aberrant mesolimbic dopamine signalling is implicated in motivational symptoms, however direct manipulations to these pathways have yielded suboptimal therapeutic effects. GPR88 is an orphan G protein-coupled receptor highly expressed in the striatum on medium spiny neurons, and therefore well-placed to modulate striatal signalling. While the phenotype of *Gpr88* knockout mice supports a disruption of motivational pathways, it is unclear whether GPR88 is involved in reward valuation and/or effort-based decision making in a sex-dependent manner, and if this involves altered dopamine function.

**Methods:** In male and female *Gpr88* knockout mice, we used touchscreen-based progressive ratio, with and without reward devaluation, and effort-related choice tasks to assess motivation and cost/benefit decision making, respectively. To explore whether these motivational behaviours were related to altered striatal dopamine, we quantified expression of dopamine-related genes and/or proteins, and used [^18^F]DOPA PET and GTPγ[^35^S] binding to assess pre- and postsynaptic dopamine function, respectively.

**Results:** We show that male and female *Gpr88* knockout mice display greater motivational drive than wild-type mice, which was maintained following reward devaluation. Further, we show that cost/benefit decision making is impaired in male, but not female, *Gpr88* knockout mice. Surprisingly, we found that *Gpr88* deletion had no effect on striatal dopamine by any of the measures assessed.

**Conclusion:** Our results highlight that GPR88 regulates motivational control of behaviour through a dopaminergic-independent mechanism, providing further support for GPR88 as target for mood symptoms in psychiatric disorders.

## Introduction

Aberrant motivational control is a common feature of psychiatric disorders with symptoms ranging from avolition and apathy to compulsive reward-seeking (1–3). Current treatments for such symptoms are often ineffective, possess unwanted side effects or, in some instances, exacerbate motivational deficits. As such, there is a significant need for novel treatments that are effective without compliance-prohibitive side effects.

The striatum is a key integrator of cognitive, motor and limbic circuitry that collectively function to regulate motivated behaviours (4). Midbrain dopaminergic projections and glutamatergic projections from numerous cortical and subcortical areas converge onto striatal medium spiny neurons (MSNs) to form cortico-striatal-thalamic loops, critical brain circuits for controlling movement, habit formation and reward processing (5). Within the striatum, functional subdivisions are associated with distinct aspects of reward learning and decision making. The dorsolateral (or sensorimotor) striatum is responsible for stimulus-response associations and habitual behaviours, while the dorsomedial (or associative) striatum is important for response-outcome associations and goal-directed behaviour (5). Finally, the ventral (or limbic) striatum is implicated in motivation and outcome evaluation. Concerted activity across all functional subdivisions—particularly with respect to dopamine signalling— is required for intact motivational control, therefore striatal targets are well positioned to modulate various aspects of motivational dysfunction.

GPR88 is an orphan G protein-coupled receptor almost exclusively expressed in the striatum, on both D_1_- and D_2_-expressing MSNs (6). GPR88 expression is altered following use of drugs of abuse, highlighting a potential role in regulating motivation and reward-related pathways (7,8). Indeed, *Gpr88* knockout mice display increased alcohol-seeking and risk-taking behaviour, and increased appetitive motivation (9,10). However, it is unclear if GPR88 is involved in reward valuation and/or effort-based decision making in a sex-dependent manner, and if behavioural changes are due to maladaptations to the dopaminergic system. To address these questions, we probed the motivational phenotype of male and female *Gpr88* knockout mice using the rodent touchscreen system, which offers a translational platform to measure cognitive behaviours with better alignment of preclinical and clinical test constructs and outcomes (11,12). We tested the effect of reward devaluation on progressive ratio breakpoint to assess the potential for altered reward valuation in *Gpr88* knockout mice. We also assessed cost/benefit decision making in *Gpr88* knockout mice in an effort-related choice task where animals were given the option of a low-effort/low-reward or high-effort/high-reward. Finally, we assessed the effect of *Gpr88* deletion on the dopamine system at a gene, protein and functional level using RT-qPCR, western blotting, [^18^F]DOPA PET imaging and GTPγ[^35^S] binding, respectively. Our work aims to clarify the role of GPR88 in motivation and reward-related behaviours and provide further validation of its utility as a target for dysfunctional motivational control in psychiatric disorders.

## Material and Methods

### Animals

*Gpr88^Cre/Cre^* mice were obtained from Jackson Laboratories (13). Wild type (WT) and *Gpr88^Cre/Cre^* mice were bred in-house from heterozygote crossing and used for all behavioural procedures, qRT-PCR, and western blot. *Gpr88* CRISPR mice (*Gpr88*^-/-^) were generated using CRISPR/Cas9 gene editing (Supplementary Methods) and then used for [^18^F]DOPA PET after being validated against behavioural data from *Gpr88^Cre/Cre^* mice (Supplementary Figure 1).

Mice had access to water and food *ad libitum* and housed in a 12-hour light/dark cycle at constant temperature and humidity. All experiments were approved by The Florey Institute of Neuroscience and Mental Health Animal Ethics Committee (16-034-FINMH, 18-132-FINMH) or by the Monash University Animal Ethics Committee (17661).

The animals used for each experiment are as follows:

Cohort 1: *Gpr88*^Cre/Cre^ mice approximately 11 weeks of age at the start of touchscreen operant training (*Gpr88^Cre/Cre^* n=12 males, n=11 females; WT littermates n=11 males, n=10 females); Progressive Ratio, Effort-Related Choice
Cohort 2: *Gpr88*^Cre/Cre^ mice approximately 11 weeks of age at the start of touchscreen operant training (*Gpr88*^Cre/Cre^ n=12 male, n=12 female; WT littermates n=12 male, n=12 female); Progressive Ratio with devaluation, RT-qPCR
Cohort 3: *Gpr88*^Cre/Cre^ mice 12-40 weeks (*Gpr88*^Cre/Cre^ 6 male, 5 female; WT littermates n=6 male, n=5 female); western blotting
Cohort 4: *Gpr88*^-/-^ mice 8-16 weeks (*Gpr88*^-/-^ n=7 male, WT littermates n=7 male); [^18^F]DOPA PET
Cohort 5: *Gpr88*^-/-^ mice 12-40 weeks (*Gpr88*^-/-^ n=6 males, n=5 females; WT n=6 males, n=4 females); striatal GTPγ[^35^S] binding

### Touchscreen apparatus

The touchscreen automated system (Campden Instruments Ltd.. Cambridge UK) was used as previously described (14,15). In brief, the system is composed of a touch-sensitive screen, a liquid reward-delivery magazine situated opposite to the touchscreen, infra-red (IR) beams and black Perspex side walls. A black Perspex mask with five response windows was placed in front of the touchscreen to limit the area that the mice were allowed to touch. The operant chamber was placed inside a sound- and light-attenuated box with house light, tone generator, IR camera and ventilating fan. Strawberry flavoured milk (3D Devondale, Australia (Experiment 1) or Nippy’s, NSW, Australia (Experiment 2)) was used as liquid reward. The system was controlled using Whiskers and Animal Behaviour Environment Test (ABET) software (Lafayette Instrument Company, Lafayette, IN, USA).

### Behavioural procedures

For all behavioural experiments, mice were housed under a reversed light-dark cycle condition to allow testing during the active phase. Mice (8 weeks of age) were acclimatised from a standard light-dark cycle to the reversed light-dark cycle for two weeks prior to any intervention. One week prior to touchscreen operant testing, mice (approx. 10 weeks of age) were weighed for three consecutive days to establish their free feeding body weight and then food restricted to reach 85% of their free feeding body weight by giving limited amounts of chow pellets every day. For two days immediately prior to the beginning of operant training, mice were exposed to a small amount of the liquid reward in their home cages to prevent neophobia. Touchscreen training and tasks protocols were adapted from (14).

#### Touchscreen operant training

Operant training was conducted as previously described (14). Briefly, to habituate mice to the testing apparatus, mice were placed in the touchscreen chamber free to explore for 20 minutes over two consecutive days. During initial touch training, a stimulus was presented (white square) for 30 seconds. Once the stimulus was turned off, a tone was issued, the reward magazine was lit and 20 μL of reward was dispensed. A 5 second inter-trial interval (ITI) followed and a new trial began. If the white square was touched while illuminated, the stimulus was switched off, the tone issued, and the magazine was lit. Triple reward delivery was issued on these trials. The session ended after the mouse consumed 30 rewards or after 60 minutes, whichever came first. The criterion to move to the next training phase was to consume 30 rewards. During the Fixed Ratio Training, mice underwent fixed ratio (FR) 1 training which consisted of 20 μL reward delivery after a single touch response to the stimulus. The session ended after completion of 30 trials or after 60 minutes. Mice then underwent FR2 (two responses required for reward delivery), FR3 (three responses required for reward delivery) and FR5 (five responses required for reward delivery; 4 days) to ensure mice established robust responding Completion criteria for FR training was 30 trials within 60 minutes.

#### Progressive Ratio (PR)

Once mice completed touchscreen operant training, they were tested on the progressive ratio schedule as previously described (14). The number of screen touches required for reward delivery now increased linearly by 4 (progressive ratio (PR) 4: 1, 5, 9, 13, 17 touches etc.) or by 8 (PR8: 1, 9, 17 touches etc.) on each trial. The session ended after 60 mins or if no screen response was made or no magazine entry was detected for 5 mins. Mice were tested on the progressive ratio schedule for four consecutive days to ensure all mice reached and maintained baseline performance.

#### Effort related choice (ERC)

Mice were tested for effort related choice using different fixed ratio schedules as previously described (14). The effort choice was evaluated by placing three pellets of standard rodent chow on the floor within the touchscreen chamber prior to testing. Mice were tested on FR 16, 32 and 5 for four consecutive sessions each. Each session finished after 60 minutes or after the completion of 30 trials. As soon as the session ended, mice were immediately removed and any residual food pellets weighted.

#### Experiment 1: progressive ratio and effort related choice

After touchscreen operant training, mice from Cohort 1 underwent progressive ratio testing (four days PR4 and four days PR8) followed by effort related choice testing.

#### Experiment 2: progressive ratio with devaluation

After touchscreen operant training, mice from Cohort 2 underwent progressive ratio training (four days PR4, baseline). Reward devaluation experiments commenced the day following session four of PR baseline testing. A baseline day was run between interventions to re-establish baseline performance and ensure there was no impact on performance for the next intervention.

#### Pre-feeding with chow

Mice were individually housed and given 30 minutes of free access to standard laboratory chow prior to the PR session. Chow was weighed before and after the 30 minutes to determine the amount of chow each mouse had consumed. This experiment spanned three consecutive days.

#### Pre-feeding with strawberry milk

Mice were individually housed and given 30 minutes of free access to strawberry milk prior to the PR session. Milk was weighed before and after the 30 minutes to determine the amount of milk each mouse had consumed. This experiment spanned three consecutive days.

#### Free feeding

Mice were then provided free access to standard laboratory chow *ad libitum* until their weights increased and stabilised. Once free feeding weights had stabilised, mice were then tested on the PR schedule with continued free access to chow for four consecutive days.

### Behavioural measures

All touchscreen data was recorded using the ABET recording software (Lafayette Instrument Co, IN, USA). For progressive ratio, the main measure of interest was the animals’ breakpoint, which is the number of touches made in the last successfully completed trial. For effort related choice, the main measure was the number of trials completed and amount of chow consumed during the testing session.

### [^18^F]DOPA PET

All scanning was performed at Monash Biomedical Imaging. Mice from Cohort 4 were used in the experiments (Supplementary Methods).

### Striatal GTPγ[^35^S] binding

Striatal membranes were prepared from Cohort 5 and GTPγ[^35^S] binding was assessed following addition of pramipexole (Supplementary Methods).

### Quantitative real time-PCR (qRT-PCR)

Brain tissue of naive and Cohort 2 mice was collected and expression of dopamine-related genes was quantified by qRT-PCR (Supplementary Methods).

### Western blotting

Striatal brain tissue from Cohort 3 was collected and expression of dopamine-related proteins was quantified by western blotting (Supplementary Methods).

### Statistical Analysis

Statistical analysis was performed using Prism version 7 or 8 (GraphPad, CA, USA). Repeated measures analysis of variance (ANOVA) and analysis of covariance (ANCOVA) were used to analyse behavioural experiments. Student’s t-test and ANOVA were used for RT-qPCR and western blot analysis. When appropriate, post hoc analysis was done using the Tukey multiple comparisons test, with a significance level set at P<0.05.

## Results

### *Gpr88* deletion increases motivation for a palatable reward

Progressive ratio tasks have been used in both animals and humans to assess a subject’s ability to maintain responding for a reward as response requirements increase. The breakpoint, or the number of responses at which the subject stops responding, provides a measure of motivation encompassing the reinforcing properties of the reward and the point at which effort outweighs the benefit of obtaining that reward. We used a touchscreen-based progressive ratio task to assess motivation in *Gpr88^Cre/Cre^* mice, where the number of touches required to elicit a strawberry milk reward increased linearly by 4 (PR4) or 8 (PR8) each trial. We found that there was no effect of sex on average breakpoint over the test sessions (Supplementary Figure 2A), therefore male and female data were combined. *Gpr88^Cre/Cre^* mice had a significantly higher breakpoint than WT littermates at both reinforcement schedules (Figure 1), indicating greater motivation for a palatable reward. This finding reflects previous reports of increased reward-seeking behaviour in *Gpr88* knockout mice (9,10).

**Figure 1.**
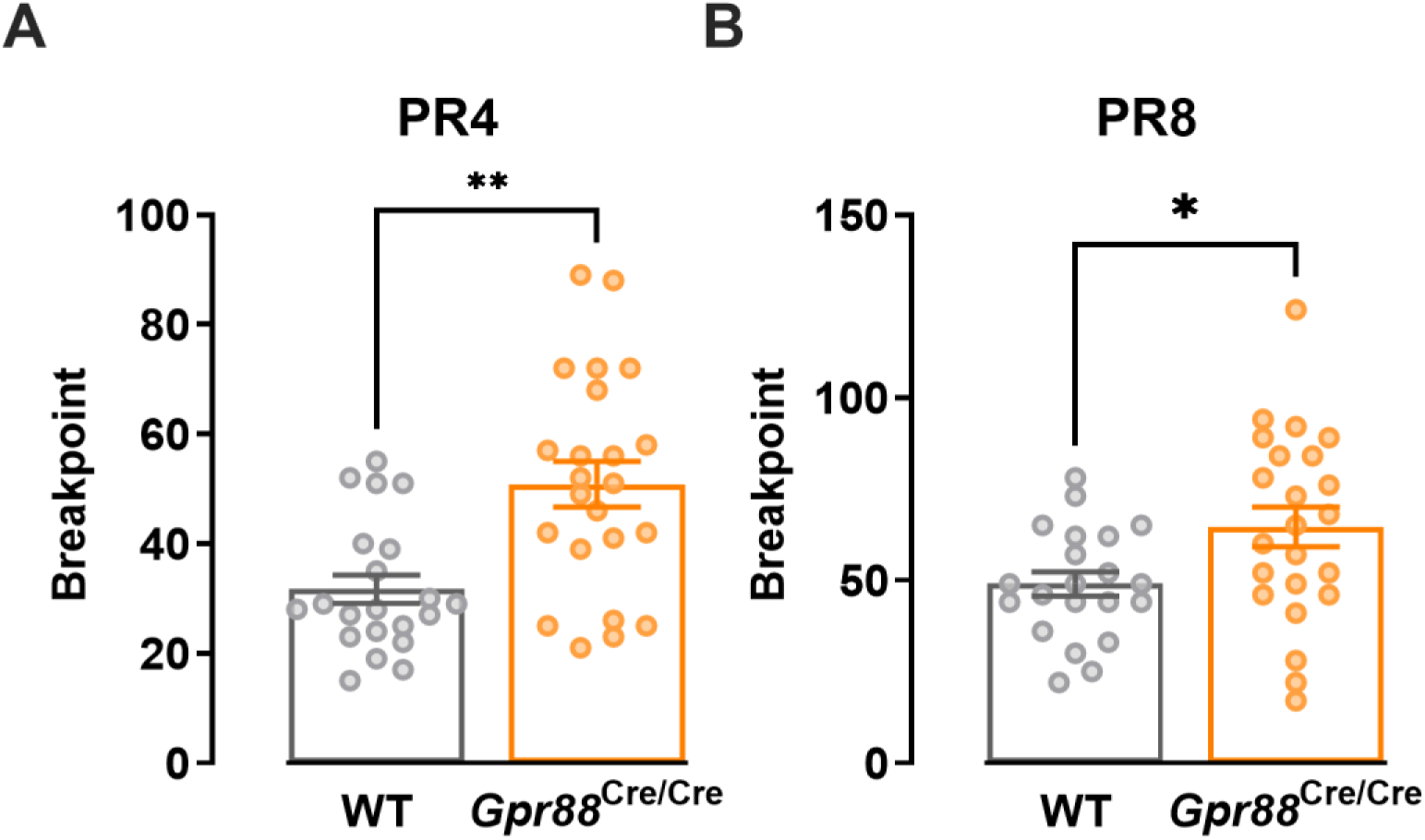
*Gpr88* deletion increases motivation towards a palatable reward at progressive ratio schedules of **(A)** 4 and **(B) 8**. **P<0.01, *P<0.05 determined by two-tailed unpaired t test; n=21-23 (WT males n=11, females n=10; *Gpr88^Cre/Cre^* males n=12, females n=11). Individual data points presented with mean ± SEM. PR4, progressive ratio 4; PR8, progressive ratio 8.

### *Gpr88*^Cre/Cre^ mice show increased motivation despite devaluation of the reward

GPR88 has a reported role in feeding and metabolism (16), therefore the increased progressive ratio breakpoint observed in *Gpr88*^Cre/Cre^ mice may be explained by a metabolic, rather than motivational, phenotype. To investigate the dependence of food intake on the progressive ratio breakpoint measure, we tested a separate cohort of *Gpr88*^Cre/Cre^ mice on PR4 following reward devaluation, where animals were given free access to either chow or milk prior to touchscreen testing, and under free-feeding conditions. Again, we found no main effect of sex on breakpoint therefore male and female data were combined (Supplementary Figure 2B). In accordance with our earlier observations, *Gpr88^Cre/Cre^* mice still displayed a significantly higher breakpoint than WT mice (Figure 2A).

**Figure 2.**
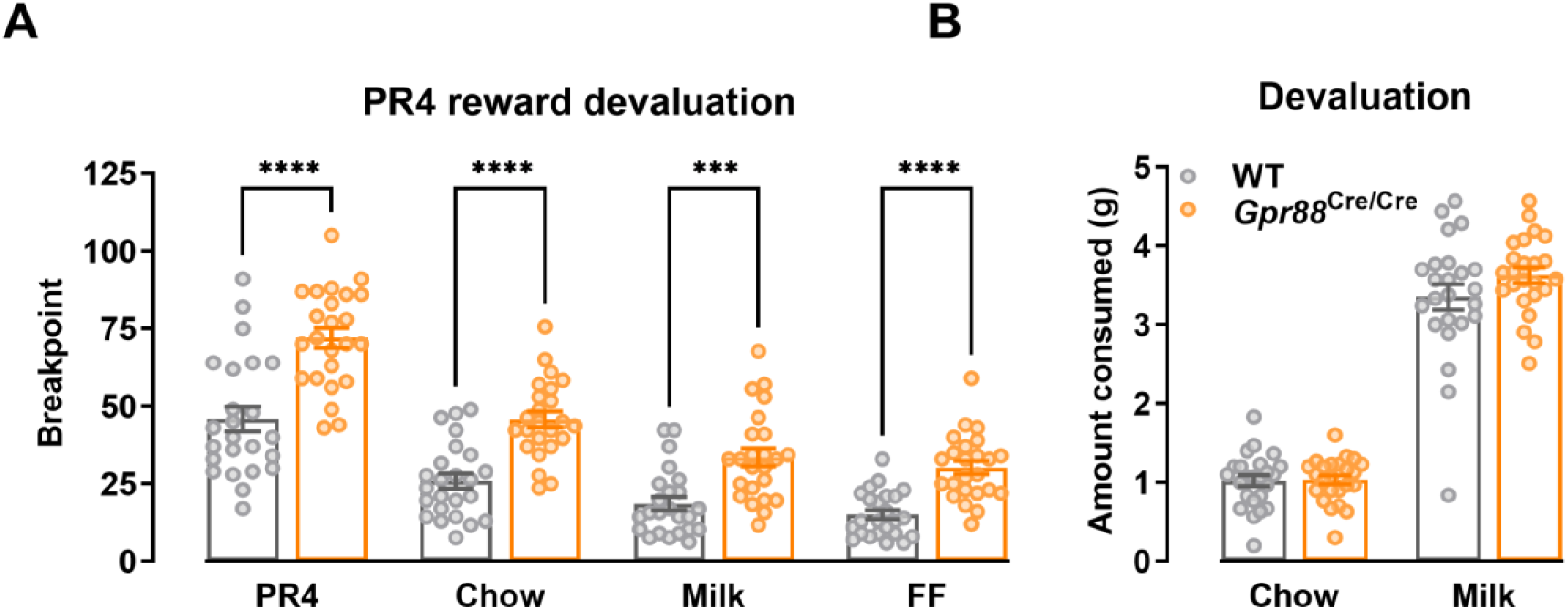
**(A)***Gpr88^Cre/Cre^* mice show increased motivation compared to WT mice following reward devaluation with chow, milk and free-feeding. RM two-way ANOVA, devaluation P<0.0001. ***P<0.001, ****P<0.0001 determined by RM two-way ANOVA with Geisser-Greenhouse correction and Šídák’s multiple comparisons test **(B)**Consumption of chow and milk prior to testing does not significantly differ between Gpr88 knockout and WT mice. RM two-way ANOVA with Šídák’s multiple comparisons test; chow P=0.997, milk P=0.133. n=24 (WT males n=12, females n=12; *Gpr88^Cre/Cre^* males n=12, females n=12). Individual data points presented with mean ± SEM. FF, free feeding.

Compared to testing under food restricted conditions, devaluation (chow and strawberry milk) or free-feeding prior to PR4 sessions, decreased breakpoint across both *Gpr88^Cre/Cre^* and WT mice as expected (Figure 2A). Despite this, *Gpr88^Cre/Cre^* mice retained a significantly higher breakpoint than WT mice in all conditions tested (Figure 2A). Importantly, there was no significant difference in the amount of chow or milk consumed by *Gpr88^Cre/Cre^* and WT mice prior to PR4 sessions (Figure 2B). Together this suggests that *Gpr88* plays a role in regulating motivational processing, and loss of GPR88 increases responding for a palatable reward independently of whether energy requirements are met, and does not impact mechanisms of satiety.

### Reward-related decision making is impaired in male, but not female, *Gpr88*^Cre/Cre^ mice

Cost/benefit analysis is a critical component that drives motivated behaviour: individuals evaluate estimated costs (i.e. effort) against the estimated value of an expected reward to optimise action selection, dysfunction of which is associated with negative symptoms of psychiatric disorders like schizophrenia (17). To next evaluate cost/benefit decision making in *Gpr88^Cre/Cre^* mice, we used a touchscreen-based effort-related choice paradigm in which animals could either make operant touches to receive the strawberry milk reward, or consume chow that was freely available. The expectation is that as the required effort to obtain the more preferred reward choice (strawberry milk) increases, animals will instead choose to consume the low effort choice (standard chow) (14). Here, we observed sex-dependent effects (Figure 3): at fixed ratio schedules of 16 (FR16) and 32 (FR32), male *Gpr88^Cre/Cre^* mice completed a greater number of trials than male WT mice, reaching significance at FR16. Interestingly, no genotype effect was observed in female mice. When the effort required for the reward was reduced to a very low level by using a FR5 schedule, all groups completed the majority of trials without significant differences between genotype or sex. As session duration varied between animals, we corrected chow consumption for time spent in the chamber. Somewhat surprisingly, despite the increased number of trials completed, male *Gpr88^Cre/Cre^* mice consumed the same amount of chow during testing as female *Gpr88^Cre/Cre^* and WT animals.

**Figure 3.**
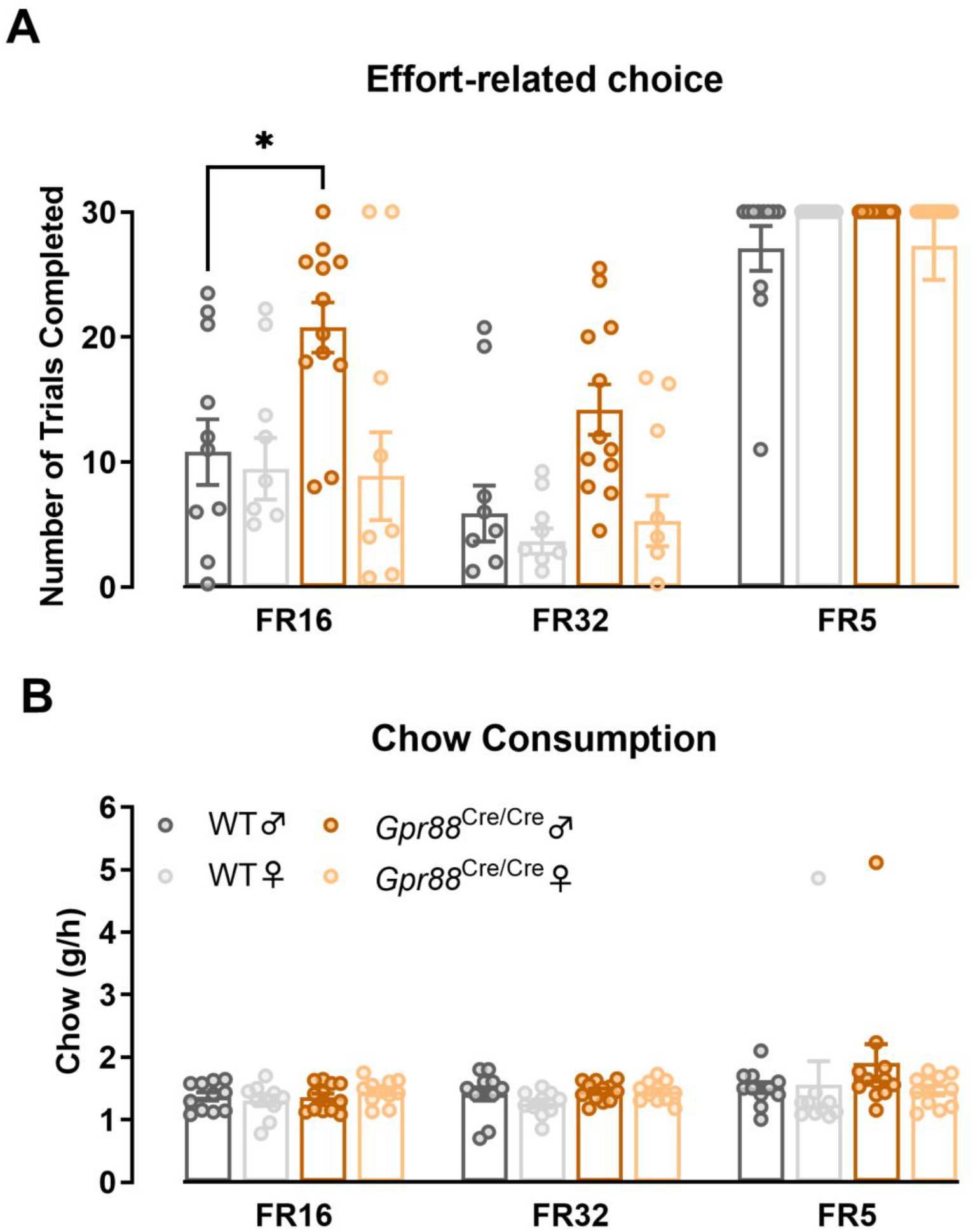
**(A)**Reward-related decision making is impaired in male, but not female, *Gpr88^Cre/Cre^* mice when effort is high, **(B)**despite equal consumption of chow across all fixed ratio schedules. *P<0.05 determined by RM two-way ANOVA with Geisser-Greenhouse correction and Tukey’s multiple comparisons test; n=10-12. Individual data points presented with mean ± SEM.

### *Gpr88* deletion does not alter mRNA or protein levels of dopamine-related targets

To determine whether the increased motivational phenotype in *Gpr88^Cre/Cre^* mice was associated with transcriptional changes to dopamine-related genes, we quantified expression of those involved in dopamine synthesis (*Th, Ddc*), signalling (*Drd1, Drd2, Drd3, Ppp1r1b*), transport (*Slc6a3, Slc18a1, Slc18a2*), and metabolism (*Comt, Maoa, Maob*) in the dorsal and ventral striatum using qRT-PCR (Table 1; Figure 4A, B).

**Figure 4.**
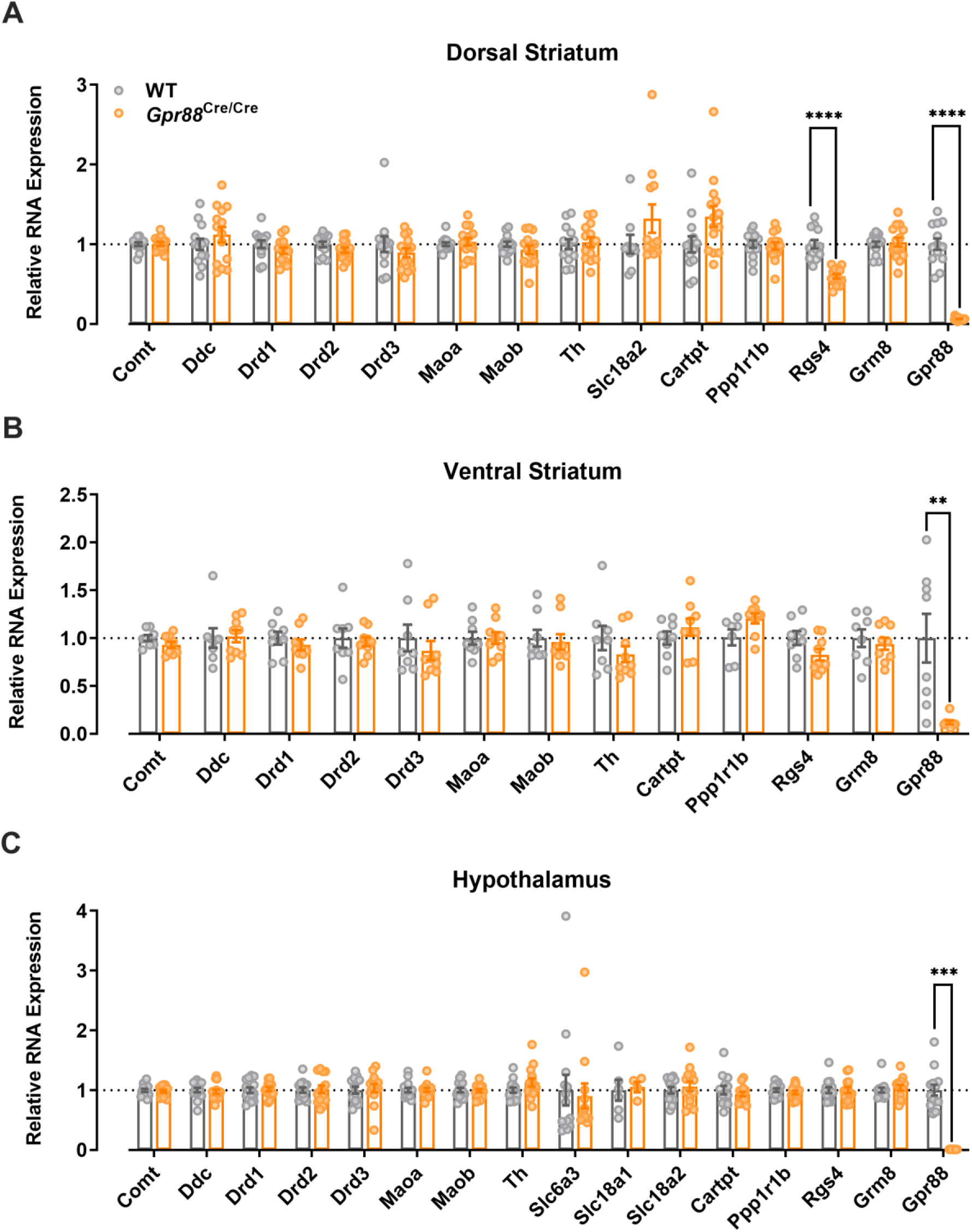
Expression of dopamine related genes is unchanged in *Gpr88^Cre/Cre^* mice in both the **(A)**dorsal and **(B)**ventral striatum, and **(C)**hypothalamus. **P<0.01, ***P<0.001, ****P<0.0001 determined by multiple Mann-Whitney tests with Holm-Šídák’s correction for multiple comparisons; dorsal striatum and hypothalamus n=13-14; ventral striatum n=8-9. Individual data points presented with mean ± SEM.

**Table 1.** Description of genes and proteins quantified by qRT-PCT and/or western blotting.

| Gene | Protein | Description |
| --- | --- | --- |
| Comt | Catechol-O-methyltransferase | Dopamine metabolism |
| Maoa | Monoamine oxidase A | Dopamine metabolism |
| Maob | Monoamine oxidase B | Dopamine metabolism |
| Drd1 | Dopamine receptor D<sub>1</sub> | Dopamine signalling |
| Drd2 | Dopamine receptor D<sub>2</sub> | Dopamine signalling |
| Drd3 | Dopamine receptor D<sub>3</sub> | Dopamine signalling |
| Ddc | L-Aromatic amino-acid decarboxylase | Dopamine synthesis |
| Th | Tyrosine hydroxylase | Dopamine synthesis |
| Slc6a3 | Dopamine transporter | Dopamine transport |
| Slc18a1 | Vesicular monoamine transporter 1 | Dopamine transport |
| Slc18a2 | Vesicular monoamine transporter 2 | Dopamine transport |
| Cartpt | Cocaine- and amphetamine-regulated transcript prepropeptide | Food intake, body weight, reward |
| Ppp1r1b | Dopamine- and cAMP-regulated phosphoprotein, Mr 32 kDa | Integrator of striatal neurotransmission |
| Rgs4 | Regulator of G protein signalling 4 | Inhibits signal transduction |
| Grm8 | Metabotropic glutamate metabotropic receptor 8 | Negative control |
| Gpr88 | G protein-coupled receptor 88 | Positive control |

In addition, we investigated genes reported to be differentially expressed in *Gpr88* knockout mice (*Cartpt, Rgs4*)(13,16) and hypothalamic tissue, where no changes in dopamine-related genes were expected (Figure 4C). *Gpr88* and *Grm8* were included as positive and negative controls, respectively. Expression of *Gpr88* was significantly reduced in both dorsal and ventral striatal regions, and the hypothalamus (Figure 4; multiple Mann-Whitney test with Holm-Šídák’s correction for multiple comparisons; dorsal striatum P<0.0001, ventral striatum P=0.007, hypothalamus P=0.0002).

No significant changes were found for any of the dopamine-related genes investigated. However, *Rgs4* was significantly lower in the dorsal, but not ventral striatum, suggesting that a previous finding of striatal Rgs4 downregulation in *Gpr88^Cre/Cre^* mice is driven by changes in the dorsal region (Figure 4; multiple Mann-Whitney test with Holm-Šídák’s correction for multiple comparisons; dorsal striatum P<0.0001, ventral striatum P=0.807; (13)).

Given that mRNA expression is not always indicative of protein expression, we further investigated enzymes and transporters involved in mesolimbic dopamine signalling in whole striatal tissue. In particular, we quantified levels of tyrosine hydroxylase (TH), amino acid decarboxylase (AADC), dopamine transporter (DAT), and monoamine oxidase A and B (MAO-A, MAO-B) by western blotting. Consistent with results from qRT-PCR, we found no changes in expression of dopamine-related proteins in *Gpr88^Cre/Cre^* mice, with respect to WT mice (Figure 5; RM two-way ANOVA, genotype P=0.948).

**Figure 5.**
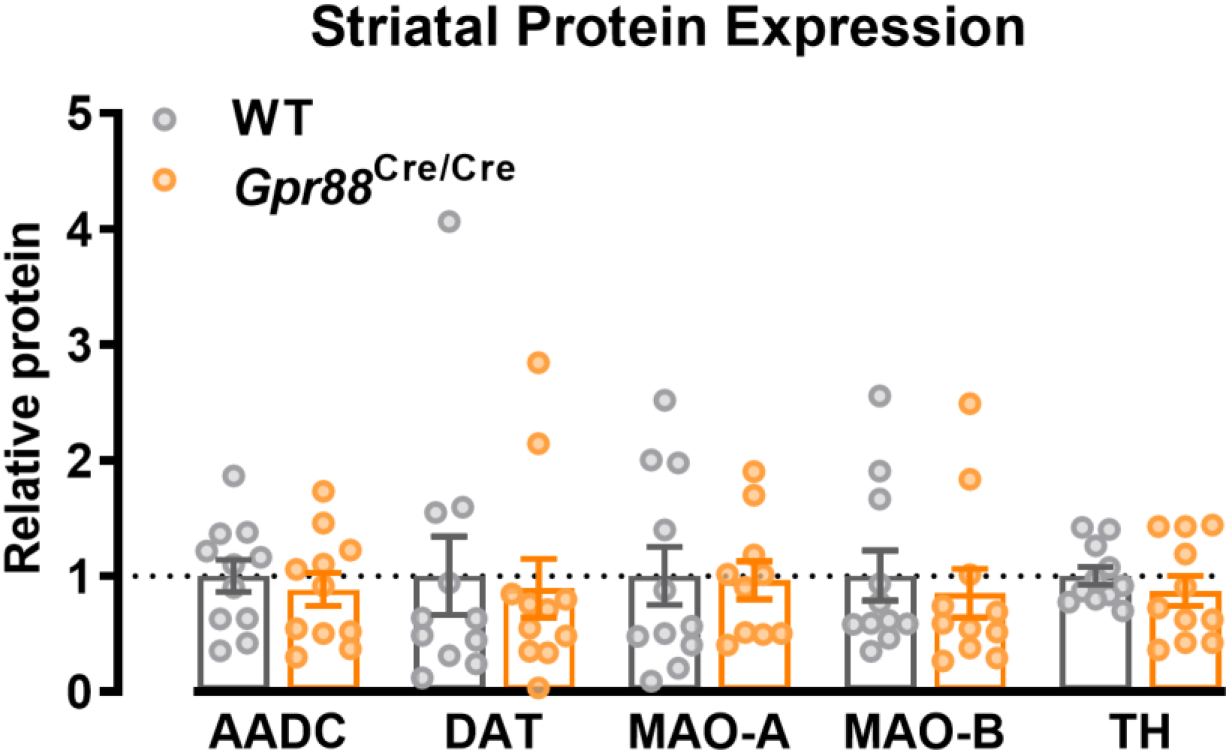
Striatal expression of dopamine-related proteins is unchanged in Gpr88^Cre/Cre^ mice (P>0.05 determined by multiple Mann-Whitney tests with Holm-Šídák’s correction for multiple comparisons; n=11). Individual data points presented with mean ± SEM.

### Striatal dopamine synthesis capacity and dopamine D_2_/D_3_ receptor function are unchanged in *Gpr88*^-/-^ mice

While no genotype-dependent differences were found in the expression of dopamine-related genes and proteins, an obvious limitation to these techniques is that they do not provide functional information. In order to address this, we investigated dopamine synthesis capacity using [^18^F]DOPA PET and dopamine D_2_/D_3_ receptor function using GTPγ[^35^S] binding.

[^18^F]DOPA PET is commonly used in clinical studies and provides a composite measure of presynaptic dopamine function. Striatum and cerebellum uptake of [^18^F]DOPA was corrected for bodyweight and radiotracer dose, and specific striatal uptake was calculated by subtracting the cerebellum time course from striatal time course (18). Specific striatal [^18^F]DOPA uptake was unchanged in *Gpr88*^-/-^ mice (Figure 6A). Similarly, dopamine synthesis capacity, indexed as K_i_^Cer^, was not significantly different between WT and *Gpr88*^-/-^ mice (Figure 6B). The K_i_^Cer^ values obtained are lower than previously reported, likely due to slight variations in experimental processes (19).

**Figure 6.**
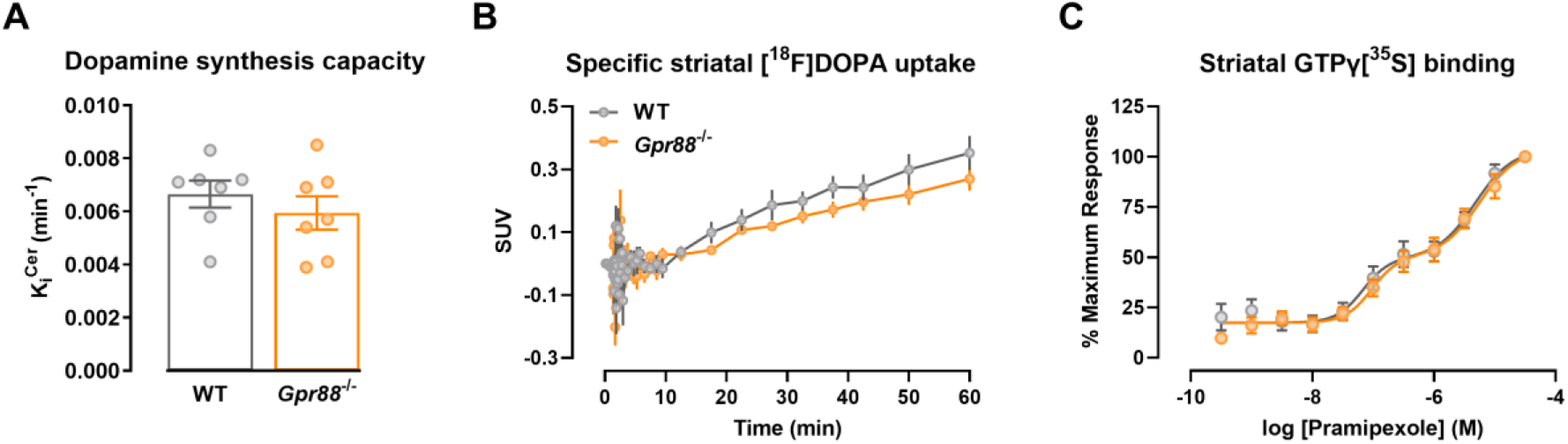
Gpr88 deletion does not affect striatal uptake of [^18^F]DOPA (RM two-way ANOVA with Geisser-Greenhouse correction, genotype P=0.439) or **(B)**striatal dopamine synthesis capacity (unmatched t-test, P=0.395); n=7). **(C)**Striatal dopamine D_2_/D_3_ receptor function is unchanged in *Gpr88*^-/-^ mice (F-test, P=0.75; n=10-11). Data presented as mean ± SEM.

Having established that presynaptic dopamine content is unchanged, we studied post-synaptic dopamine D_2_/D_3_ receptor function in striatal membranes prepared from WT and *Gpr88*^-/-^ mice by GTPγ[^35^S] binding. We found there was no effect of sex on GTPγ[^35^S] binding, therefore male and female data were combined. The dopamine D_2_/D_3_ receptor agonist, pramipexole, stimulated GTPγ[^35^S] binding in a concentration-dependent and biphasic manner with potencies for the two phases of approximately 100 nM and 5 μM in membranes from both WT and *Gpr88*^-/-^ mice (likely reflecting multiple Gα_i/o_ coupled receptor subtypes being activated by pramipexole in the native preparation; Figure 6C). Notably, there was no significant effect of genotype on pramipexole potencies.

## Discussion

In this study, we investigated the effect of *Gpr88* deletion on motivational behaviour and any associated changes to striatal dopamine function. We report that male and female *Gpr88*^Cre/Cre^ mice display increased motivation for a palatable reward, which is maintained following reward devaluation and occurs independently of food restriction. Interestingly, we found that *Gpr88* deletion affects cost/benefit decision making in a sex-dependent manner, whereby male, but not female, *Gpr88*^Cre/Cre^ mice display a high-effort bias. Finally, we investigated changes to dopamine function at a gene, protein and functional level but, somewhat surprisingly, found no differences between *Gpr88* knockout and WT mice. Together this work further delineates the role of GPR88 in motivation and reward-related pathways, and highlights it regulates these behaviours via a putative non-dopaminergic mechanism for motivational control.

Dopamine signalling is heavily implicated in reward and motivation-related pathways. In progressive ratio tasks, both D_1_ and D_2_ antagonists reduce breakpoint, while inhibiting dopamine reuptake or increasing dopamine release increases breakpoint (20–22). Despite the clear effect of *Gpr88* deletion on increasing the breakpoint in the progressive ratio task, we did not identify any changes to striatal dopamine at a gene, protein or functional level. Notwithstanding, there is some evidence to suggest that GPR88 deletion may indirectly potentiate dopamine signalling. First, *Gpr88* deletion increases excitability of both D_1_ and D_2_ GABAergic MSNs, which account for ~95% of the striatal neuronal population (13). MSNs form reciprocal connections with midbrain dopaminergic neurons, and project via the direct D_1_-expressing striatonigral pathway, or the indirect D_2_-expressing striatopallidal pathway (4,23). These pathways are embedded in broader cortico-striatal-thalamo-cortical loops that regulate a range of behaviours, including motivation (24,25). While activity of dopaminergic projections to the striatum may remain unchanged in *Gpr88* knockout mice, supported by normal dopamine synthesis capacity (Figure 6B) and postsynaptic dopamine receptor function (Figure 6C), it is possible that increased excitability of MSNs leads to downstream potentiation of dopamine signalling.

Second, we confirm previous reports that *Gpr88* deletion downregulates striatal expression of RGS4, and found this was specific to the dorsal striatum (Figure 4A)(13). RGS proteins are critical regulators of GPCR function, effectively switching off signalling following receptor activation (26). Notably, expression of RGS4 is upregulated following dopamine D_2_, but not D_1_, receptor activation, suggesting a selective role of RGS4 in the inactivation of D_2_ signalling pathways (27). As such, *Gpr88* deletion may increase the magnitude and duration of D_2_ receptor signalling, leading to an imbalance in D_1_ *vs* D_2_ receptor activation. Given that both D_1_ and D_2_ receptor antagonists reduce breakpoint in progressive ratio tasks, it is likely that a balance in signalling, rather than signalling at either receptor *per se*, is critical for the maintenance of normal motivation (20). This balance of dopamine signalling required for normal function is not exclusive to motivation, but has been shown for a number of behavioural outputs including working memory and locomotor activity (28,29). Indeed, *Gpr88* knockout animals show increased sensitivity to the locomotor-inducing effects of amphetamine, providing further support for an imbalance of striatal dopamine signalling (30).

While the progressive ratio task provides a basic measure of reward valuation, the effort-related choice task provides better insight into cost/benefit decision making given the competing choice between a low effort/low reward or high effort/high reward. In this task, male, but not female, *Gpr88^Cre/Cre^* mice had a higher breakpoint than WT mice indicating a high effort/high reward bias (Figure 2A). Increased synaptic dopamine and antagonism of the adenosine A2A receptor have been shown to shift preference towards the high effort reward (22,31,32). Interestingly, these pharmacological manipulations simultaneously decrease intake of the low effort reward, whereas male *Gpr88*^Cre/Cre^ mice consume the same amount of chow as wildtype animals (Figure 2B). The mechanisms underlying the sex-dependent impairment of cost/benefit decision making in *Gpr88*^Cre/Cre^ mice is largely unclear, but may be related to the metabolic phenotype of *Gpr88* knockout mice - with which males display a more pronounced phenotype (16).

An important consideration in our work, and related approaches, is that we used free-operant behavioural paradigms of reinforcement learning that involve food restriction, which affects motivation in its own right (Figure 2A). In the case of *Gpr88* knockout mice, the interplay between motivation and energy requirements is of particular interest as GPR88 is expressed in the hypothalamus and has an established role in feeding, body composition and energy expenditure (16). Lau et al. report that *Gpr88* knockout mice have reduced spontaneous food intake and energy expenditure, with respect to WT mice, without changes to body weight gain or physical activity (16). Surprisingly, we observed no significant differences in food intake between *Gpr88*^Cre/Cre^ and WT mice (Figure 2B; 3B), but found that female, but not male, *Gpr88*^Cre/Cre^ mice had a consistently lower body weight than WT mice both before and during behavioural procedures (Supplementary Figure 3). Together with reports of reduced body weight in male *Gpr88* knockout mice, this suggests genotype effects on weight may be sensitive to environmental factors (33). Indeed, *Gpr88* deletion reportedly increases fasting-induced food intake under a high-fat, but not normal chow diet, highlighting a complex role of GPR88 in the maintenance of energy homeostasis. We found that the increased motivation in *Gpr88* knockout mice occurred independently of food intake, however it is unclear exactly how the combination of food restriction and strawberry milk reinforcement interacts with energy homeostasis in *Gpr88* knockout mice, and therefore how it may influence appetitive motivation.

In conclusion, this study confirms that *Gpr88* deletion alters motivational responding for a palatable reward, and provides evidence that this effect is not driven by the metabolic phenotype previously reported in *Gpr88*^-/-^ mice. Furthermore, we show a sex-depedent effect of *Gpr88* deletion on effort-related decision making, where male, but not female, mice make more high-effort/high-reward choices than WT mice. Given that both tasks are sensitive to manipulation by dopaminergic drugs, we hypothesised that changes in motivation may be driven by underlying changes to dopamine function. Somewhat surprisingly, we found no dopamine-related changes at a gene, protein, or functional level suggesting motivational regulation occurs independently of dopamine signalling.

## Acknowledgments

This work was supported by a National Health and Medical Research Council Project Grant (1104371) to CL and JN. JN was supported by an Australian Research Council Future Fellowship (140101327). The authors acknowledge the scientific and technical assistance of the National Imaging Facility, a National Collaborative Research Infrastructure Strategy (NCRIS) capability, and Monash Biomedical Imaging, Monash University, Ms Yao Lu, Monash Institute of Pharmaceutical Science, Monash University, and Dr Robyn M. Brown, Florey Institute of Neuroscience and Mental Health. The mutant animals were produced via CRISPR genome editing by the Monash Genome Modification Platform (MGMP), Monash University as a node of Phenomics Australia. Phenomics Australia is supported by the Australian Government Department of Education through the National Collaborative Research Infrastructure Strategy, the Super Science Initiative and the Collaborative Research Infrastructure Scheme.

