## Supplementary Material for "*Gpr88* deletion impacts motivational control independently of striatal dopamine function"

### Supplementary Methods

#### *Gpr88*$^{-/-}$ CRISPR mice

*Gpr88* CRISPR mice were generated using CRISPR/Cas9 gene editing using sgRNA1 5′ AAAGAACAGGAGAACGGGGG 3′ sgRNA2 5′ AGCCCCAGGGAAGATCCCAG 3′. CRISPR-Cas9 RNAs (crRNA) were ordered from Integrated DNA Technologies (IDT) and incubated with transactivating CRISPR RNA (tracrRNA) to form a functional guide RNA complex. Cas9 nuclease was purchased from IDT (IDT Alt-R® S.p. HiFi Cas9 Nuclease V3) and incubated with the guide RNAs to form a ribonucleoprotein (RNP) complex. The ssDNA repair template was generated using the Guide-it Long ssDNA Production System (Takara) according to the manufacturer's instructions. Cas9 nuclease (30ng/ml), gRNAs 1 & 2 (30ng/ml) & ssDNA repair template (30ng/ml) were microinjected into the pronucleus of C57BL/6J zygotes at the pronuclei stage. Injected zygotes were transferred into the uterus of pseudo pregnant F1 females.

#### [$^{18}$F]DOPA PET

30-45 minutes prior to [$^{18}$F]DOPA treatment, mice were dosed with peripherally-restricted inhibitors of L-DOPA metabolism, benserazide (formulated in MilliQ $H_2O$ at 10mg/kg in a 5 ml/kg dose volume, i.p.) and entacapone (formulated in 20% (2-hydroxypropyl)-β-cyclodextrin in $dH_2O$ pH=4-5 at 40 mg/kg in 5 ml/kg dose volume, i.p). Prior to scanning, mice were anaesthetised with isoflurane in air and a tail-vein cannula was inserted. Animals were then placed in the μPET-CT scanner (Siemens, Germany) and a 60-minute scan commenced. After 1 minute of baseline reads, mice were dosed with [$^{18}$F]DOPA (4.31 ± 0.85 MBq). At the end of the PET scan, a 10-minute CT scan commenced for attenuation correction and to provide structural references. Following this, animals were transferred to the MRI scanner (Bruker, USA) to acquire a structural MRI using FLASH with TR/TE = 60/8 ms, resolution = 0.156 x

0.156 x 0.156 $mm^3$ (FOV = 14.976 x 14.976 x 7.8 $mm^3$, matrix size = 96 x 96 x 50), slice thickness = 7.8 mm, slice number = 1, average = 4.

PET data was histogrammed into 47 frames (4 x 20s, 10 x 3s, 14 x 5s, 6 x 30s, 4 x 60s, 7 x 300s, 2 x 600s), reconstructed using filtered back projection and corrected for CT attenuation, radiotracer decay and deadtime. Using Inveon Research Workspace software (Siemens, Germany), structural MRI was registered to PET and CT data and used to define 3D ROIs of the striatum and cerebellum, which was used as a reference region in lieu of arterial input.

Time activity curves (TACs) for the striatum and cerebellum were generated and modelled by Patlak graphical analysis. Striatal $K_i^{Cer}$, the influx rate constant, was calculated from a linear regression of data between 10-60 minutes relative to the cerebellum. $K_i^{Cer}$ provides a composite measure of striatal dopamine synthesis, storage, release and metabolism.

**Striatal GTPγ[$^{35}$S] binding**

Striatal membranes were prepared from tissue dissected from adult *Gpr88*$^{-/-}$ mice (12-40 weeks of age, males and females). In brief, striatal tissue was kept on ice and homogenised using a hand held homogeniser (Polytron PT1200E, ThermoFisher Scientific, Waltham, Massachusetts, USA) in an iced-cold buffer containing 20mM HEPES and 10mM EDTA and pH 7.4. Homogenised tissue was then centrifuged at 500g for 5 min at 4 ℃ (Heraeus Multifuge 3SR+ Centrifuge, ThermoFisher Scientific, Waltham, Massachusetts, USA) the pellet was homogenised and centrifuged again as above. The resulting supernatant was then centrifuged at 40,000g for 1h at 4 ℃ (Sorvall Evolution RC, ThermoFisher Scientific, Waltham, Massachusetts, USA). The pellet was then resuspended in an iced cold buffer containing 20mM HEPES, 1mM EDTA, pH 7.4 and passed through a 30G needle. Protein concentration was determined using a BCA assay (Thermo Fisher Scientific, CA, USA) following manufacturer instructions. GTPγ[$^{35}$S] binding was assessed using 20 µg of protein per well. In brief, membrane preparations were incubated with and without ligands (Pramipexole Dihydrochloride (Sigma #1598), at concentrations from 30 µM to 300 pM, diluted in assay buffer, and Haloperidol (Sigma #1512), at a single concentration of 10 nM, diluted in 1% DMSO in assay buffer) for 1h at 37 °C in assay buffer (25 mM HEPES, 100 mM NaCl, 5 mM MgCl2, 2 mM CaCl2, 0.2 mM EGTA, 0.01% Pluronic F127 and 20 µg/mL saponin, pH 7.4) containing 100 µM GDP and 0.2 nM GTPγ[$^{35}$S]. The reaction was stopped by rapid filtration through Whatman GF/C filters, which were immediately washed with an ice-cold wash buffer (0.9% NaCl), dried and dissolved in Microscint™-0, and counted using a MicroBeta$^{2}$® 2450 Microplate Counter (PerkinElmer, Waltham, Massachusetts, USA). Basal binding was assumed to be the specific GTPγ[$^{35}$S] binding in absence of agonist.

**Quantitative real time-PCR (qRT-PCR)**

Brain tissue was collected from ventral striatum (WT male n=4, female n=4; *Gpr88*$^{Cre/Cre}$ male n=4, female n=5), dorsal striatum (WT male n=5, female n=8; *Gpr88*$^{Cre/Cre}$ male n=6, female n=8), and hypothalamus (WT male n=6, female n=8; *Gpr88*$^{Cre/Cre}$ male n=6, female n=8) and quickly frozen in dry ice. RNA was extracted from each sample using the Bioline RNA ISOLATE II RNA mini kit (Bioline, London, UK) following the manufacturer instructions. cDNA was synthesised using the tetro-cDNA synthesis kit (Bioline, London, UK) following the manufacturer instructions. The qRT-PCR was performed using LightCycler 480 SYBR green (Roche, Basel, Switzerland) and the Bio-Rad CFX384 Touch Real-Time PCR Detection System (Bio-Rad, CA, USA). CFX Manager Software (Bio-Rad, CA, USA) was used to analyse RT-qPCR data and normalise gene expression to house-keeping genes by employing the DCt method where the average triplicate Ct value for each sample was subtracted. *Gpr88*$^{Cre/Cre}$ DCt values were then normalised to WT samples for each gene of interest to obtain the DDCt. β-actin and GAPDH were used as reference house-keeping genes (see list of genes and primer used in Table 1).

**Western blotting**

Mice from Cohort 3 were humanely killed by cervical dislocation and heads were decapitated, the brain was then removed. For the dopamine pathway-related proteins the striatum was dissected and immediately placed in dry ice. Tissue was sonicated in lysis buffer containing 5mM HEPES, 300mM NaCl, 1% NP40, 10% glycerol, 0.4% DDM, 1:100 Halt(™) protease and phosphatase inhibitors (Thermo Fisher Scientific, CA, USA) and made up to volume with MilliQ water. A BCA assay (Thermo Fisher Scientific, CA, USA) was performed on the lysed tissue to determine protein concentration in each sample. Samples were loaded into 8-16% Mini-PROTEAN TGX Stain-Free Protein Gels (Bio-Rad, CA, USA). Precision Plus Protein Dual Color Standards (Bio-Rad, CA, USA) were used as molecular weight markers. Proteins were then electrophoretically transferred to polyvinylidene difluoride membranes (Immobilon, Millipore Corp., MA, USA) overnight at 4 °C and 30 V. Membranes were blocked for one hour at room temperature in 5% BSA in 1x PBST and then probed with primary antibodies overnight at 4 °C (Table 2). Blots were washed at room temperature three times for 5 minutes each in blocking buffer. Blots were then incubated with secondary antibody for 1 hour at room temperature. Membranes that were probed with HRP-conjugated secondary antibody were incubated for 5 minutes with Pierce enhanced chemiluminescence (ECL) Plus substrate (Thermo Fisher Scientific, CA, USA) before imaging. Blots were visualised via fluorescence using the Amersham Typhoon (GE Healthcare Bio-Sciences Corp, MA, USA) or via chemiluminescence using the Bio-Rad Chemidoc Imaging System (Bio-Rad, CA, USA). Images were analysed using Image J via densitometry by normalising the bands of interest with housekeeping proteins neuronal nuclear protein (NeuN) or β-actin.

### Supplementary Figures

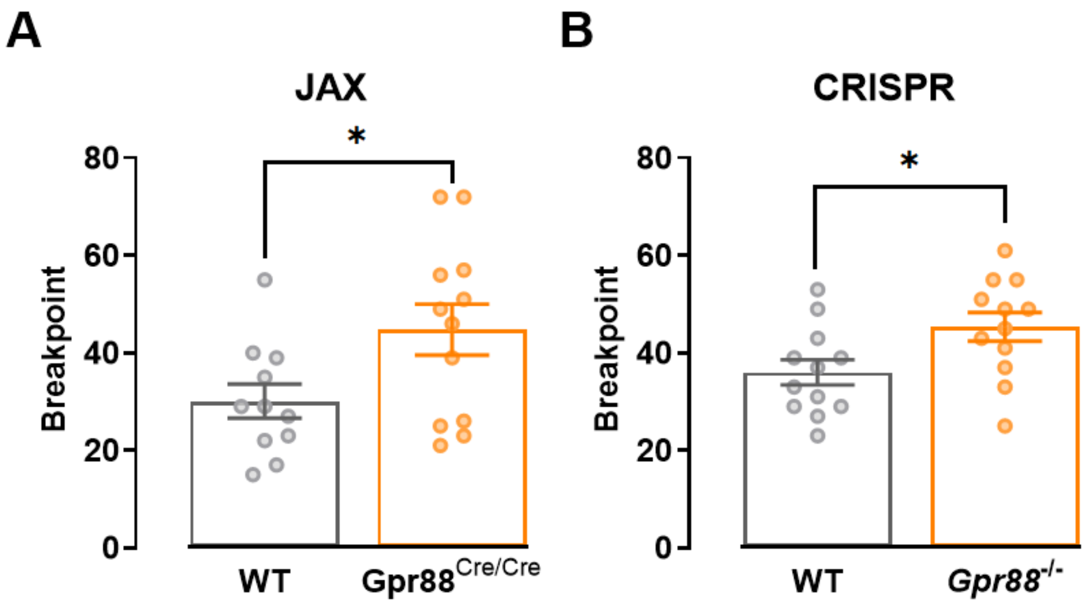


**Figure S1.** Progressive ratio breakpoint is increased in male **(A)** JAX *Gpr88*$^{Cre/Cre}$ and **(B)** CRISPR *Gpr88*$^{-/-}$ mice. *P<0.05 determined by unpaired t test; n=10-12. Individual data points presented with mean ± SEM.

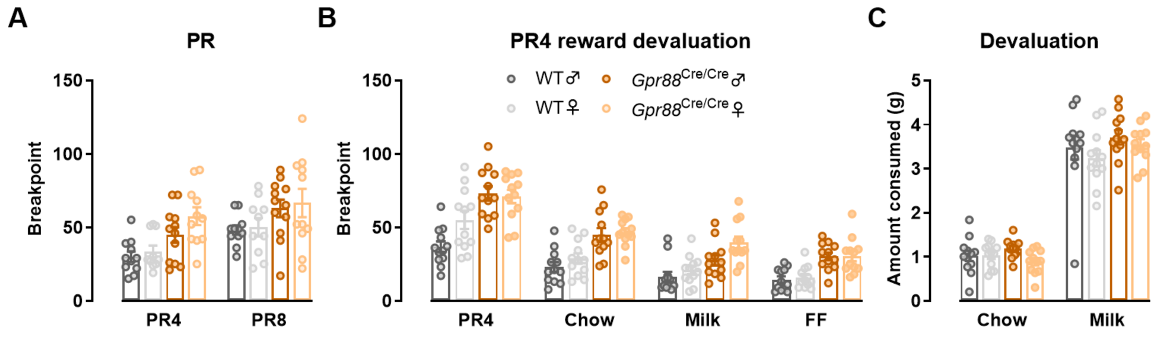


**Figure S2.** No sex-dependent effects of *Gpr88* deletion are found on **(A)** progressive ratio breakpoint at PR4 or PR8 (RM three-way ANOVA, genotype x sex P=0.5112), **(B)** PR4 breakpoint following reward devaluation (RM three-way ANOVA, genotype x sex P=0.4386) or **(C)** reward consumption during devaluation (RM three-way ANOVA, genotype x sex P=0.6412). Individual data points presented with mean ± SEM; n=10-12.

**Figure S3**

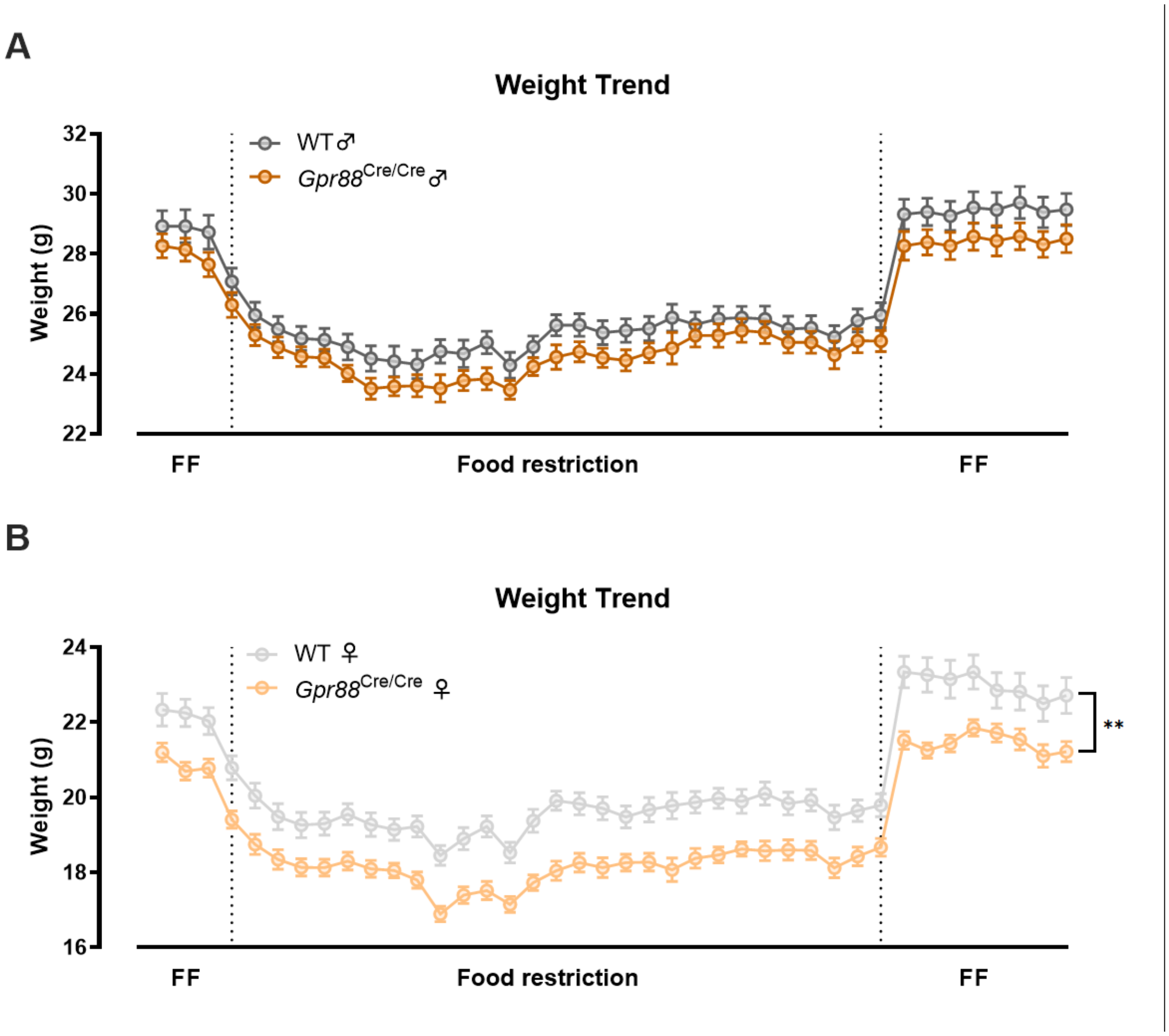


**Figure S3. (A)** Female *Gpr88*$^{Cre/Cre}$ mice have significantly lower body weight than WT mice across free-feeding and food restriction conditions in Experiment 2 (RM two-way ANOVA, genotype P=0.0010), while **(B)** male *Gpr88*$^{Cre/Cre}$ and WT mice do not significantly differ (RM two-way ANOVA, genotype P=0.1514). Individual data points presented with mean ± SEM; n=12.
